# Cisplatin resistant lung adenocarcinoma cells exhibit increased proangiogenic capacity in a microphysiological model of tumor neovascularization

**DOI:** 10.1101/2025.08.18.670623

**Authors:** Elisabet A. Olsen, G. Wills Kpeli, Omar M.K. Ahmad, Mark J. Mondrinos

## Abstract

Carcinomas commonly recur and progress rapidly after a period of remission following platinum-based therapy. This clinical scenario suggests that surviving drug-resistant tumor cells are dormant or slow cycling before re-entering a rapid growth phase. Remodeling of the recurrent tumor microenvironment (TME) contributes to high rates of metastasis, but little is known about differences in TME remodeling before therapy and after recurrence. This study explores selection for cisplatin-resistant subpopulations of A549 lung adenocarcinoma cells in culture to derive populations for modeling features of the recurrent TME. A cisplatin dose of 25 μM killed approximately 80% of the cells while sparing enough cells to allow re-expansion of sufficient cell numbers for downstream experimentation. Expanded cisplatin-resistant derivatives (Cis-R A549) exhibited features of mesenchymal transition (EMT) such as cellular hypertrophy, loss of cell-cell contacts, and upregulation of alpha smooth muscle actin mRNA. In 3D culture, Cis-R A549 spheroids were loosely aggregated and dysmorphic in comparison to the compact and spherical parent A549 spheroids. The Ki67 index of Cis-R A549 in 2D and 3D spheroid culture was markedly lower than parent A549, suggesting a state of pseudo-dormancy with slow cycling. Cis-R A549 upregulated multiple genes associated with the evolution of a more aggressive TME and displayed significantly increased proangiogenic capacity in a microphysiological model of tumor angiogenesis. This study establishes a methodological framework for engineering the recurrent TME with drug-resistant cancer cell line derivatives selected via high-dose exposure in culture. Increased angiogenesis induced by Cis-R A549 suggests that anti-angiogenic therapy may be more beneficial in the setting of recurrent disease following first-line therapies.

## Introduction

Lung cancer (LC) is the leading cause of cancer deaths in both women and men worldwide [1, 2]. LC outcomes remain poor with cumulative 5-year survival rates around 25% for all sub-types [3, 4]. The treatment history of non-small cell lung cancer (NSCLC), which comprises ∼85% of LC diagnoses, has evolved considerably over the last decade [5]. New classes of agents, such as immunotherapies and biomarker-guided targeted therapies have gained traction in clinical use [6]. These new classes of agents are used as first-line therapies for stage IV disease and are considered for earlier stages, yet the platinum-based alkylating agents such as cisplatin and carboplatin have remained a recommended component of neoadjuvant therapy for stages I-III NSCLC and for first-line adjuvant [5, 6]. These platinum-based agents cause DNA damage, thereby inducing apoptosis, inhibiting growth and proliferation, and often leading to a period of non-detectable disease when combined with surgical resection of the bulk tumor [7]. However, many factors such as stochastic heterogeneity of drug exposure in the tumor, DNA repair mechanisms, and increased expression of drug efflux pumps all contribute to a spectrum of intrinsic and acquired resistance to platinum-based therapy [8, 9]. Drug resistant tumor cells that survive therapy, more recently termed ‘persister cells’, exhibit slow cell cycling and are believed to drive eventual recurrence of more aggressive and rapidly progressing disease through unknown mechanisms [10–12].

The long history of cisplatin therapy for LC has been plagued by both intrinsic and acquired resistance [13]. Consequently, recurrence commonly occurs on varying time scales depending on the pathologic response to cisplatin therapy, although more recent trials of cisplatin combined with compounds such as have shown improved outcomes [14, 15]. While the clinical pattern of recurrence is well-established, there remains a poorly understood sequence of events beginning with selection pressure imposed by platinum-based therapy [16, 17]. This selection pressure is believed to favor the persistence of preexisting cancer stem cells (CSC), but it’s possible that adaptation to drug-induced damage can result in the acquisition of CSC phenotypes by more differentiated sub-clones [18]. Depending on the clinical scenario, events occurring on the level of surviving persister cells correspond to a period of disease dormancy that precedes recurrence and rapid metastasis [19–22]. Rapid metastasis following recurrence suggests the acquisition of more motile tumor cell phenotypes and microenvironmental changes that facilitate dissemination following persister cell re-entry into a growth phase. Angiogenesis in the recurrent TME seems to be a contributing factor along with tumor cell intrinsic properties in the sequence of recurrence and rapid progression associated with drug resistance [23–26]. Models of persister cell-induced angiogenesis are needed to accelerate the discovery of driver mechanisms and biomarkers.

Cisplatin resistance has been studied extensively in cultured lung cancer cells to define the cell biological mechanisms by which tumor cell subpopulations either evade or survive the cytotoxic effects [27–30]. There is a relative paucity of literature connecting resistant phenotypes with TME remodeling, due in part to a lack of adaptable model systems. The emergence of microphysiological systems (MPS) for cancer modeling, i.e., tumor-on-a-chip platforms and other permutations of tumor organoid-integrated culture systems, offers new opportunities for human-relevant investigation of cancer biology in vitro [31, 32]. Increased research and development of new approach methodologies (NAMs) such as MPS for cancer modeling has enabled the study of processes including tumor vascularization, immune cell recruitment, and tumor cell metastasis [33–35]. While the process of tumor angiogenesis has been modeled using MPS in various contexts [33, 36–38], none of those systems were used to model the effect of treatment history on tumor cell proangiogenic capacity. This study explores selecting for cisplatin resistance in expansion cultures of lung adenocarcinoma cells to produce derivatives for engineering models of the recurrent, drug-resistant TME with increased metrics of aggressivity. Experiments were designed to test the hypotheses that, compared to a parent line of A549 lung adenocarcinoma cells, cisplatin-resistant A549 derivatives would exhibit features of epithelial-to-mesenchymal transition (EMT), increase expression of progression-associated genes (e.g., VEGFA, TGFB1, EGFR1, IL8, ASMA), and produce 3D spheroids with a dysmorphic architecture and increased proangiogenic capacity in a MPS model.

## Results and Discussion

### Generating cisplatin-resistant A549 derivatives

Tumor recurrence following treatment is a leading cause of death in LC and other types of carcinomas, but little is known about what features of the recurrent tumor microenvironment (TME) might promote increased aggressivity and metastasis (**Figure 1A**) [39–41]. Exposing a parent line of cancer cells or patient-derived organoids established from initial biopsy to first-line therapies in vitro is a tractable approach to developing before-and-after models for defining specific TME features induced by resistance to a given compound. The current study compared cisplatin-resistant (Cis-R) A549 non-small cell lung adenocarcinoma cells with the parent line in 2D culture, 3D spheroids, and a microphysiological model of tumor angiogenesis (**Figure 1B**). A549 cells in 2D well-plate culture were treated with varying doses of cisplatin for 96 hours followed by a period of culture in maintenance medium without cisplatin to allow for recovery and expansion of surviving cells (**Figure 1C-E**).

**Figure 1:**
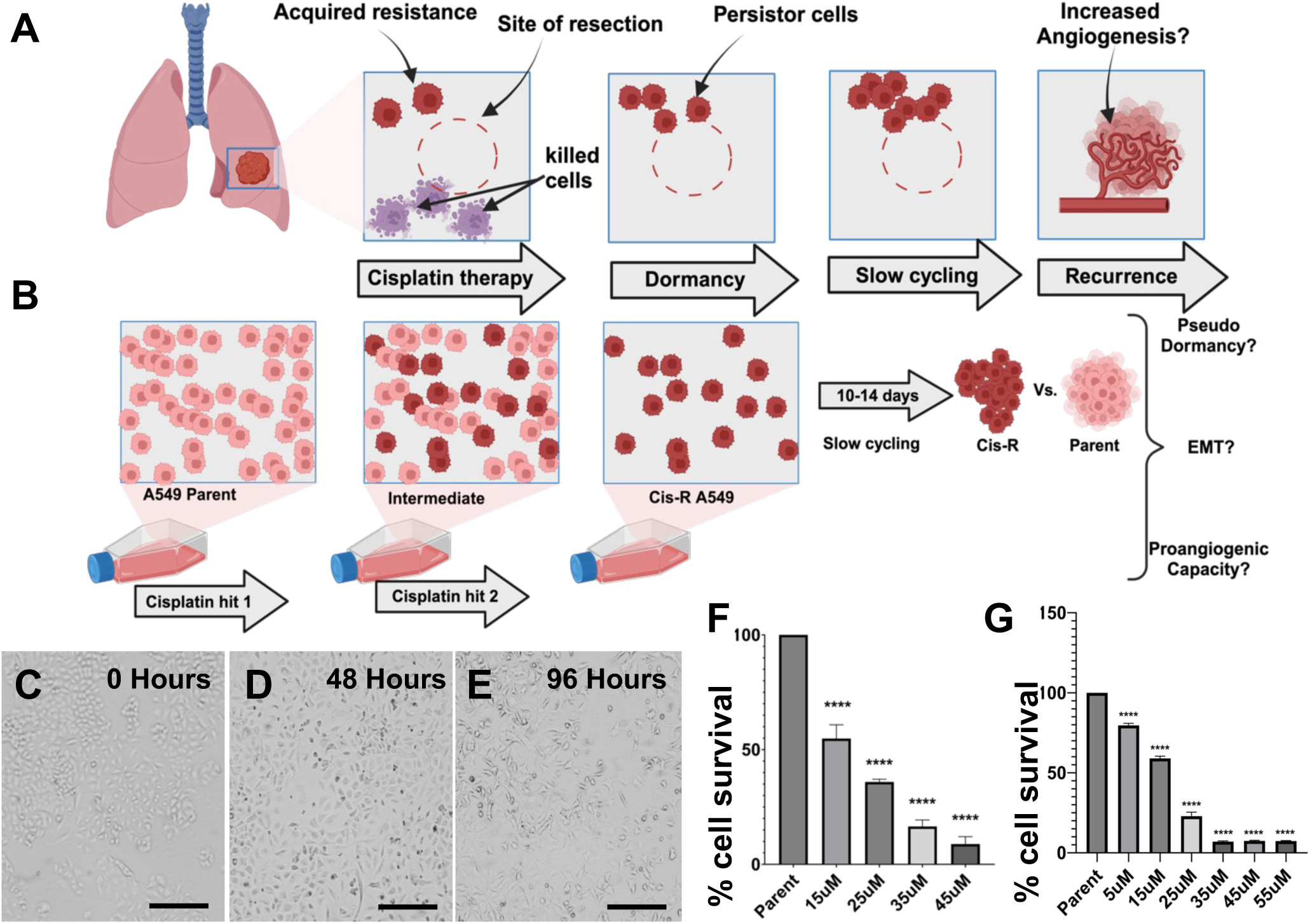
Modeling drug-resistant persister cells via selection in 2D culture. **A:** Illustration of a clinical scenario in which first-line cisplatin therapy following surgical resection kills some residual tumor cells at the surgical margins, while other tumor cells survive based on either intrinsic or acquired mechanisms of resistance. After a period of dormancy or slow cycling, these persister cells can seed the formation of local or distant recurring tumors that often progress more rapidly, perhaps due to more aggressive remodeling of the TME induced by drug resistant cells. **B:** Schematic of 2D culture protocol to select for cisplatin resistant persister A549 lung adenocarcinoma cells to test for functional changes such as altered cell cycle activity, mesenchymal transition, and increased proangiogenic capacity. **C-E:** Phase contrast micrographs of a representative selection culture at 0 hours (first cisplatin hit), 48 hours (second cisplatin hit), and 96 hours. Scale bar = 100μm. **F and G:** Percentage of surviving cells after the 96 hour 2-hit protocol at the indicated doses of cisplatin as determined by flow cytometric analysis (**F**) and the crystal violet assay (**G**). Statistical analysis was performed using a one-way ANOVA with Dunnett’s multiple comparison in GraphPad Prism v10.3.1. N = 3 independent experiments. Statistical significance is indicated by * = p < 0.05, ** = p < 0.01, *** = p < 0.001, **** = p < 0.0001. Error bars represent Standard error of the mean (SEM).

The ‘maximum recoverable dose’ was defined as a highly cytotoxic dose that spares a fraction of cells that are capable of re-expansion. Two independent assays were used to quantify dose-dependent cisplatin cytotoxicity. The crystal violet assay provides an indirect measure of cell number based on the assumptions that cell volume and per cell uptake of the dye are relatively constant, while flow cytometric analysis based on Hoechst dye labeling along with forward and side scatter provides a direct quantitative measure of viable cell numbers [42, 43]. These independent assays yielded similar results when representing the data as the percentage of live cells (**Figure 1F and 1G**). Testing a dose range of 5-55 μM revealed that doses of 5 and 15 μM killed less than 25% and 50% of the cells, respectively, while 25 μM killed approximately 70-80% of the cells. Higher doses of 35, 45, and 55 μM killed approximately 90% or more of the cells. Cells that survived the 25 μM dosing were capable of re-expansion after a 10 to 14-day period of pseudo-dormancy in maintenance medium without cisplatin, while the surviving fraction of cells dosed with 35-55 μM did not recover. Thus, 25 μM was established as the maximum recoverable dose and used for all downstream experimentation.

### Cisplatin-resistant A549 derivatives proliferate slowly and exhibit mesenchymal characteristics

Cell cycle status determined by Ki67 staining and morphometric analysis were used to quantify phenotypic differences of Cis-R A549 compared to the parent line. Parent A549 formed monolayers with discernable epithelial morphology and tight cell packing (**Figure 1C and 2A**). Cis-R A549 were qualitatively enlarged, more loosely packed, and exhibited mesenchymal-like morphology (**Figure 1E and 2B**). Notably, the Ki67 index dropped from greater than 70% in parent A549 cultures to less than 5% in recovered Cis-R A549 cultures (**Figure 2C**). This low fraction of cells with current or recent cell cycle participation explains the 10- to 14-day period required to generate re-expanded cultures suitable for passaging and downstream use. Morphometric analysis revealed significant hypertrophy in Cis-R A549 derivatives. Average cell diameters when approximating a circular shape increased from about 30 μm for the parent A549 line to 80 μm for Cis-R A549 (**Figure 2D**). Cell areas based on directly traced outlines were calculated to provide a more accurate measure of increased cell size inferred from increased spreading. Cell area increased 6-fold from approximately 1000 μm^2^ for the parent A549 line to 6000 μm^2^ for Cis-R A549 (**Figure 2E**). Nuclear area followed a similar trend, doubling from approximately 175 μm^2^ to 350 μm^2^ (**Figure 2F**).

**Figure 2:**
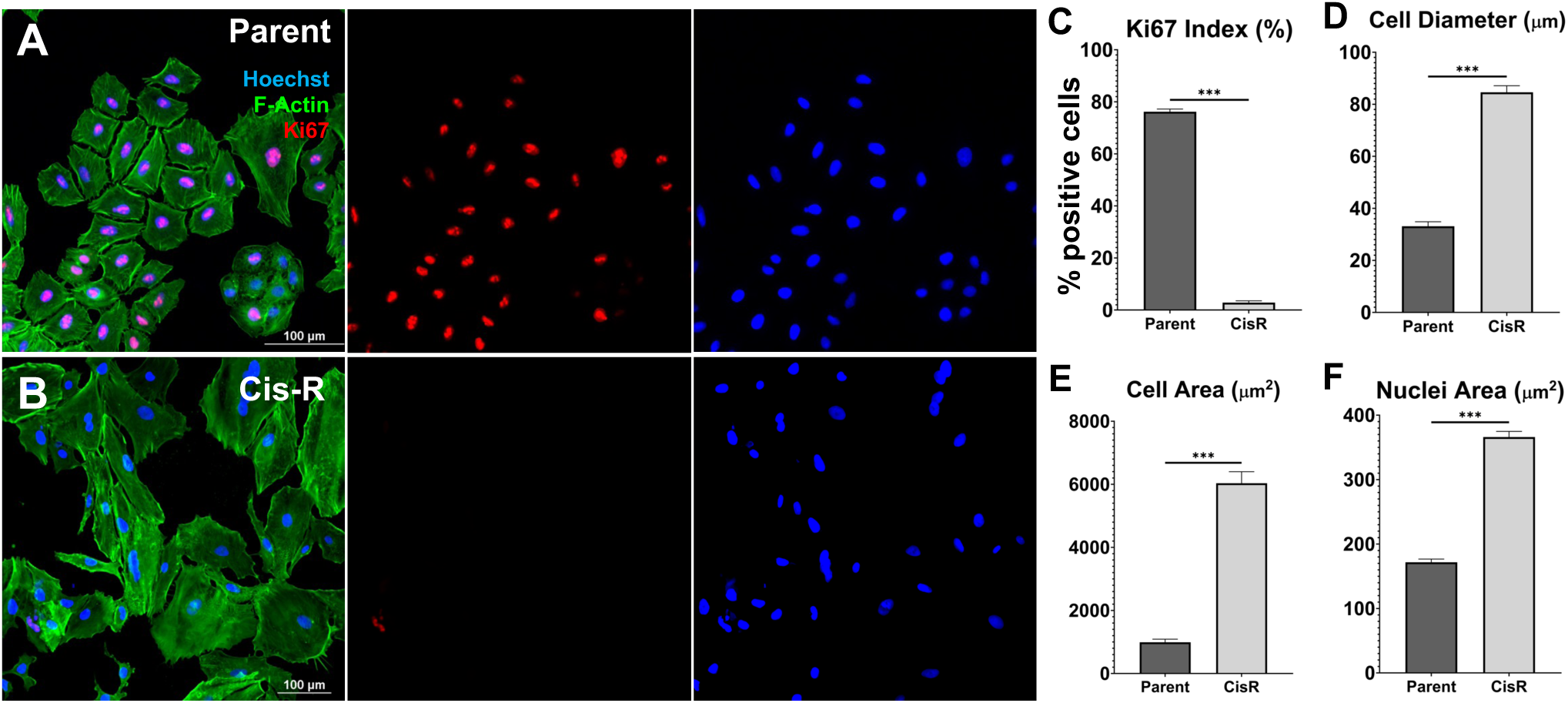
Morphological changes of A549 cells persisting after the 96 hour 2-hit cisplatin protocol at 25 μM. **A and B:** Parent A549 cells (**A**) and persister cells that survive the 2 hits with 25 μM cisplatin hits (Cis-R, **B**) immunostained for Ki67 (cell cycle marker, red) with counterstains for f-actin (phalloidins, green) and nuclei (Hoechst, blue). Scale bars = 100 μm. **C:** Parent and Cis-R Ki67 indexes defined as the percentage of Ki67 positive cells. **D-F:** Parent and Cis-R cell diameters in μm (**D**) cell areas in μm^2^ (**E**), and nuclei areas (**F**). Bar graphs are represented as the mean ± SEM, and statistical analysis was conducted in GraphPad v10.3.1 prism using unpaired t-tests with Welch’s corrections when applicable. N = 3 independent experiments. Statistical significance is indicated by * = p < 0.05, ** = p < 0.01, *** = p < 0.001

Recovered and passaged Cis-R A549 were used to generate 3D spheroids for characterization and quantitative comparison with parent A549 spheroids. Parent A549 spheroids were uniformly spherical with a compact structure indicative of tight cellular cohesion (**Figure 3A**). In contrast, Cis-R A549 formed loosely aggregated dysmorphic spheroids with gaps and cavities not seen in the parent spheroids (**Figure 3B**). Cis-R A549 exhibited a similar state of pseudo-dormancy in 3D spheroids with a Ki67 index below 5% (**Figure 3C**). By comparison, the Ki67 index of spheroids composed of parent A549 was significantly higher at 23 +/−11% (P < 0.01) (**Figure 3C**). Morphometric analysis revealed that the loosely aggregated nature of Cis-R A549 spheroids is represented by a significant decrease in solidity (**Figure 3D**). Compactness, a measure of perimeter relative to area that increases with border irregularity, was significantly increased in Cis-R A549 spheroids (**Figure 3E**). The significantly decreased form factor and circularity of Cis-R A549 spheroids also indicate transition to a dysmorphic 3D architecture (**Figure 3F and 3G**). The image area of parent A549 and Cis-R A549 spheroids were statistically equivalent, suggesting that morphometric differences are not an artifact of differing spheroid sizes (**Figure 3H**).

**Figure 3:**
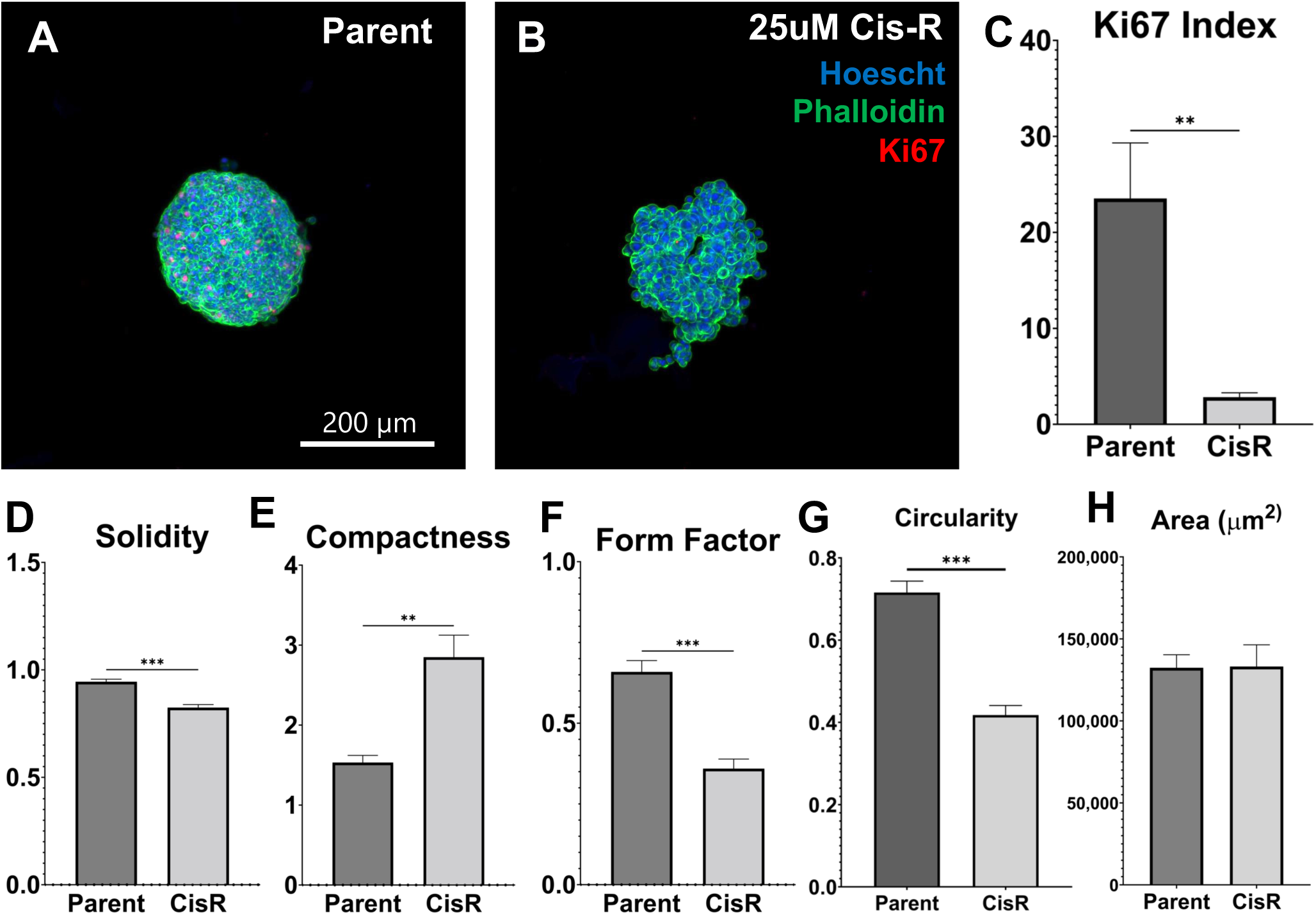
Morphology and growth characteristics of Cis-R spheroids formed after the 10–14-day recovery period in 2D culture. **A and B:** Parent A549 spheroids (**A**) and CisR persister spheroids (**B**) immunostained for Ki67 (cell cycle marker, red) with counterstains for f-actin (phalloidins, green) and nuclei (Hoechst, blue). Scale = 200 μm. **C:** Parent and Cis-R spheroid Ki67 indexes defined as the percentage of Ki67 positive cells. ** indicates P<0.01. **D-H:** Morphometric analysis of parent and Cis-R spheroid solidity (**D**), compactness (**E**), form factor (**F**), circularity (**G**), and 2D projected areas in μm^2^ (**H**). and nuclei areas (F). N = 3 independent experiments. Statistical analysis was performed using an unpaired t-test with Welch’s correction in GraphPad Prism v10.3.1. Statistical significance is indicated by * = p < 0.05, ** = p < 0.01, *** = p < 0.001. Error bars represent SEM.

### Cisplatin-resistant A549 derivatives upregulate progression-associated genes

The expression of selected genes associated with an aggressive TME were analyzed by RT-qPCR in 2D and 3D spheroid cultures of parent A549 and Cis-R A549 derivatives (**Figure 4A and B**). Epidermal growth factor receptor-1 (EGFR1) expression levels have been correlated with LC aggressivity [44, 45]. Cis-R A549 expressed approximately 3.4 +/− 2.1-fold higher levels of EGFR1 mRNA compared to parent A549 in 2D culture (not significant, ns) and 6.9 +/− 2.5-fold higher in 3D spheroids (ns) (**Figure 4A and B**). Vascular endothelial growth factor A (VEGF-A) is a multifaceted driver of tumor vascularization [46]. VEGF-A mRNA levels in parent and Cis-R A549 were nearly equivalent in 2D culture (0.8 +/− 0.2, ns), while there was a 3.7 +/− 1.5-fold upregulation in Cis-R A549 spheroids (ns). Transforming growth factor β (TGFB1) exerts pleiotropic effects in the TME that drive LC metastasis [47, 48]. While 2D cultures of Cis-R A549 slightly upregulated TGFB1 (1.6 +/− 0.6, ns), expression was significantly upregulated in Cis-R A549 spheroids (4.3 +/− 0.6, P < 0.01). Elevated expression of interleukin 8 (IL8) is associated with a higher risk of death in LC and other cancers [49]. IL8 can act as an autocrine driver of lung cancer cell growth and promotes EMT [50]. Cis-R A549 expressed 6.3 +/− 6.8-fold higher levels of IL8 mRNA relative to parent A549 in 2D, but this difference was not significant due to the high variance. By contrast, IL8 mRNA was reproducibly and significantly elevated by 4.4 +/− 0.3-fold in Cis-R A549 spheroids (P < 0.0001).

**Figure 4:**
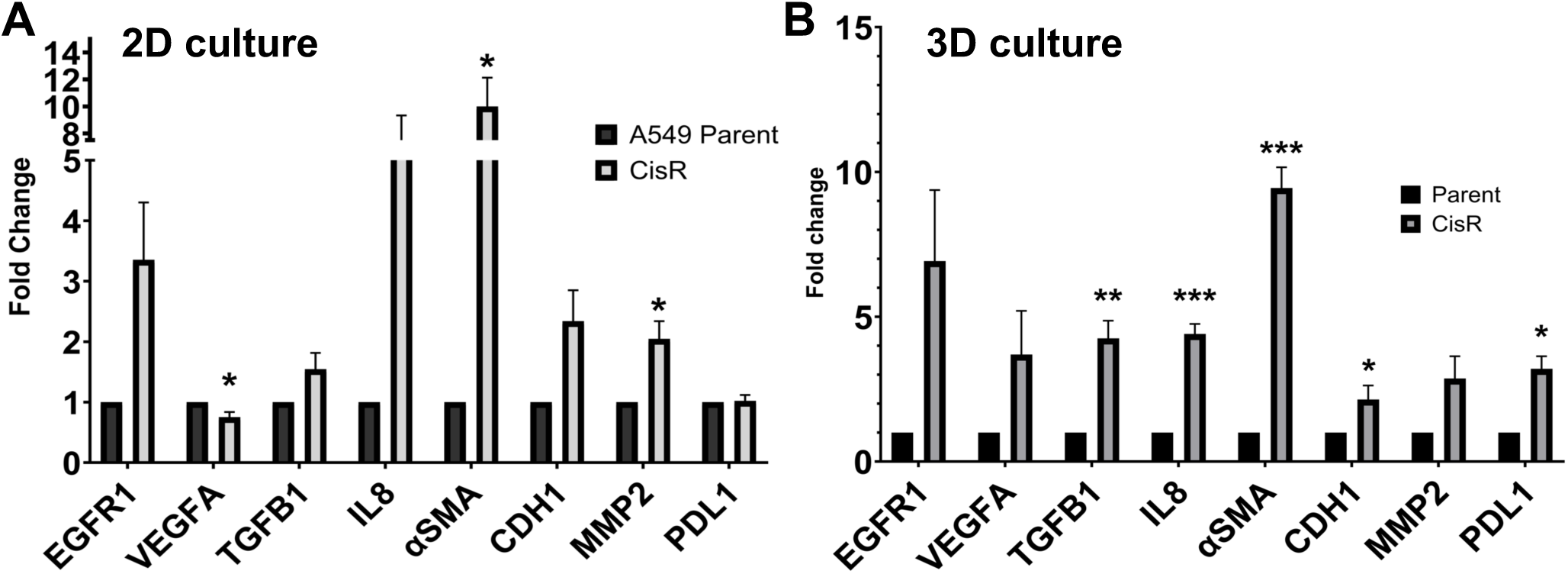
Cis-R A549 upregulate progression-associated genes in 2D and 3D culture. **A and B:** RT-qPCR analysis of EGFR1, VEGFA, TGFB1, IL8, ASMA, CDH1, MMP2, and PDL1 mRNA levels in 2D cultures **(A**) and 3D spheroids (**B**). Data are expressed as fold change of expression by Cis-R relative to parent A549. N= 3 independent experiments. Statistical analysis was performed using an unpaired t-test with Welch’s correction in GraphPad Prism v10.3.1. Statistical significance is indicated by * = p < 0.05, ** = p < 0.01, *** = p < 0.001. Error bars represent SEM.

Alpha-smooth muscle actin (ASMA) is a marker of activated contractile mesenchymal cells, the expression of which in lung tumor cells is correlated with distant metastasis and poor prognosis [51, 52]. Cis-R A549 express 10 +/− 4.8-fold higher levels of ASMA mRNA compared to parent A549 in 2D culture (P < 0.05), and 9.5 +/− 0.7-fold higher in 3D spheroids (P < 0.001). E-cadherin (CDH1) was upregulated by approximately 2-fold by Cis-R A549 in 2D culture (2.3 +/− 1.2, ns) and 3D spheroids (2.1 +/− 0.5, P<0.05). Increased expression of the type IV collagenase MMP2 is correlated with lung cancer malignancy [53, 54]. MMP2 expression was upregulated by Cis-R A549 in 2D culture (2.1 +/− 0.7, P < 0.05) and 3D spheroids (2.9 +/− 1, ns). Now a key target of immune checkpoint therapy for NSCLC, over-expression of programmed death ligand 1 (PDL1) in multiple subtypes is a known mechanism of immunological evasion via suppression of cytotoxic T cell activity [55, 56]. While CisR A549 expression of PDL1 was unchanged relative to parent in 2D cultures (1.0 +/− 0.2, ns), PDL1 expression was increased by 3.2 +/− 0.4-fold in Cis-R A549 spheroids (P<0.05).

Collectively, these data revealed important differences between 2D and 3D cultures. Only ASMA and MMP2 were significantly upregulated in 2D cultures of Cis-R A549 (**Figure 4A**). By comparison, 5 of the 8 genes assayed—TGFB1, IL8, CDH1, ASMA, and PDL1—were significantly upregulated in 3D spheroids of Cis-R A549 (**Figure 4B**). Comparing Cis-A549 gene expression in 2D vs. 3D revealed significant upregulation of TGFB1 and PDL1 for Cis-R A549 in 3D culture (**Supplemental Figure 1**). This comparison corroborates other reports of increased expression of PDL1 in 3D multicellular aggregates of carcinoma cells that highlight the importance of 3D culture systems for modeling immune evasion and evaluating immune checkpoint inhibitors [57, 58]. Similarly, modeling tumor processes driven by increased TGF-β expression and their pharmacological modulation in 3D culture is likely to increase the relevance of preclinical studies [59]. More generally, the increased expression of multiple genes by Cis-R A549 in both 2D and 3D demonstrates that cisplatin exposure altered the population-wide gene expression response to the same culture conditions experienced by the parent line. Upregulation of genes associated with poor prognosis in LC by Cis-R A549 supports the notion that drug-resistant carcinoma cell derivatives can be used for modeling more aggressive phenotypes and advanced stages of progression.

### Cisplatin-resistant A549 derivatives are more proangiogenic

Microphysiological systems (MPS) comprised of organoids, organ chips and related tissue-engineered technologies can increase the relevance of studying cancer biology in vitro by offering functional readouts of complex processes including angiogenesis and metastasis[60]. We developed an adaptable manufacturing workflow that uses inexpensive stereolithography 3D printing to fabricate resin molds for polydimethylsiloxane (PDMS) soft lithography (**Figure 5A and B**) [61]. Molded PDMS device layers are then bonded to create organ chips that are subsequently loaded with cells, hydrogels, and culture medium (**Figure 5C**). Previously, we developed a MPS assay for quantifying directional sprouting angiogenesis induced by triple negative breast cancer spheroids [62]. The MPS angiogenesis assay is performed in a membrane-free organ chip device that patterns two adjacent and contiguous layers of tissues. One layer is a vascularized ‘margin tissue’, and the adjacent layer contains tumor spheroids with or without other cell types (**Figure 5D**). This platform was used to compare sprouting angiogenesis induced by parent A549 and Cis-R A549 spheroids. Differences in spheroid morphology in hydrogels used within the device mirrored standard spheroid cultures in low attachment plates, with Cis-A549 displaying a more dysmorphic architecture and mesenchymal-like cell elongation (**Supplemental Figure 2**).

**Figure 5:**
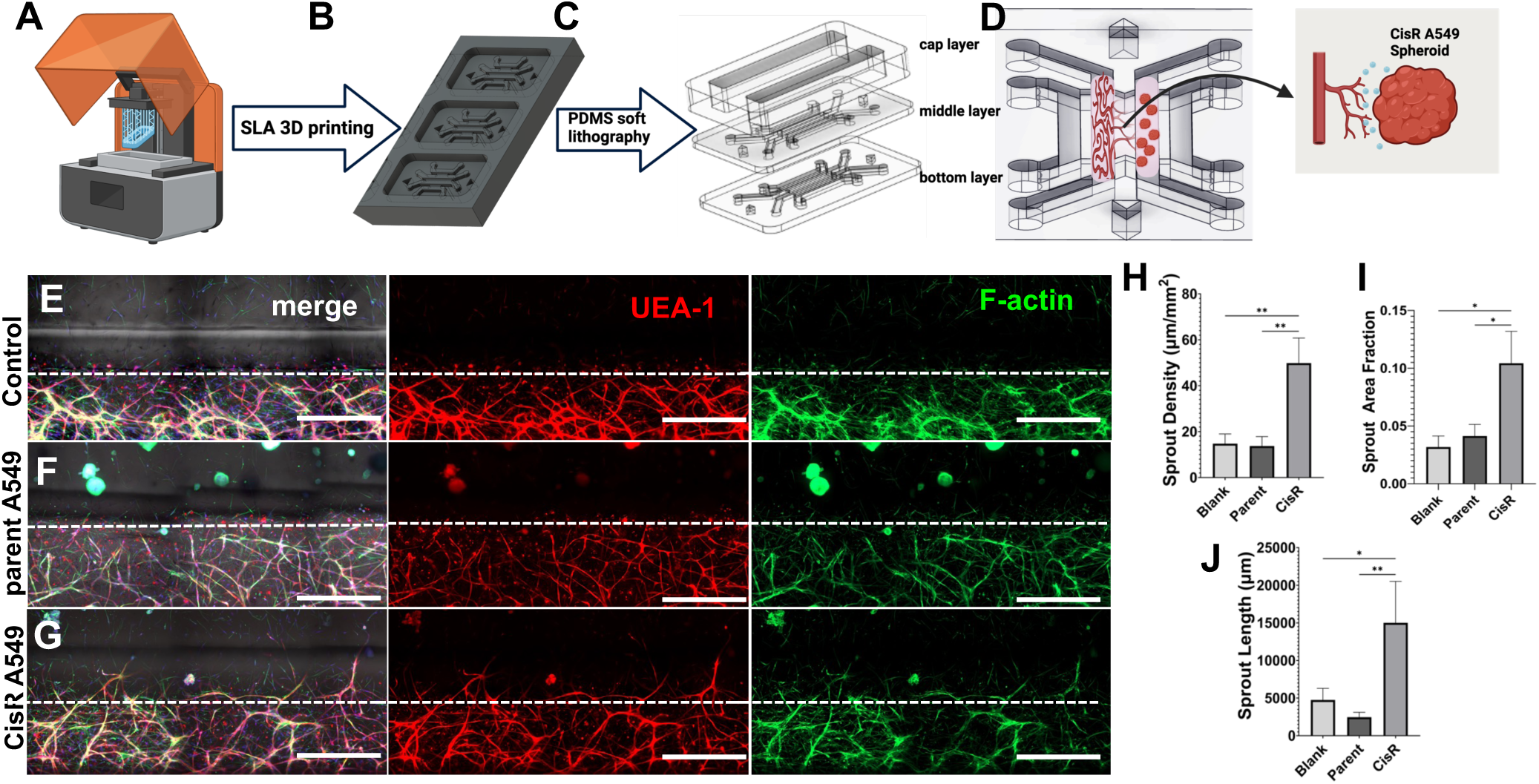
Cis-R A549 spheroids. **A-C:** Device fabrication for the MPS angiogenesis assay. A stereolithography (SLA) 3D printer (**A**) is used to fabricate resin molds (**B**) that are post-processed for use in downstream polydimethylsiloxane (PDMS) soft lithography (**C**) to produce device layers that are bonded and prepared for culture as described in Methods. **D:** The culture device patterns two adjacent and contiguous layers of tissue. A vascular margin layer (left) provides the established vasculature that sprouts into an adjacent tissue layer (right) that is kept acellular (control) or loaded with parent or Cis-R A549 spheroids. **E-G:** Endpoint whole-mount staining of devices with UEA-1 lectin (endothelial-specific, red) and Alexa488 phalloidins (F-actin, green) after 10 days of culture. Dashed lines indicate the interface between tissue layers, across which sprouting of new vessels occurs. Scale = 500μm. **H-J:** Morphometric analysis of sprout vessels. Sprout vessel densities in μm/mm^2^, p < 0.0023. (**H**), sprout area fractions p < 0.014. (**I**), and total sprout vessel lengths in μm p < 0.0023. (**J**) were significantly increased in devices with Cis-R A549 spheroids. N = 3 independent experiments. Test for significance was analyzed via one-way ANOVA with Tukey’s multiple comparison in GraphPad Prism v10.3.1. * = p < 0.05, ** = p < 0.01, *** = p < 0.001. Error bars represent SEM.

Increased expression of VEGF-A mRNA in cis-A549 spheroids (**Figure 4**) supported the hypothesis that Cis-R A549 will significantly increase sprouting angiogenesis in the MPS assay. After 10 days of culture, robust sprouting induced by coculture with Cis-R A549 spheroids (**Figure 5G**) was qualitatively apparent in comparison to control devices and coculture with parent A549 spheroids (**Figure 5E and F**). Metrics of angiogenesis including sprout length, sprout density, and x-y area covered by newly sprouted vessels were significantly increased in the Cis-R A549 devices in comparison to both other groups (**Figure 5H-J)**. Minimal sprouting in both control devices and devices containing parent A549 spheroids suggests that the parent line is not inherently angiogenic. Thus, increased potentiation of angiogenesis is likely an ability gained through phenotypic changes associated with cisplatin resistance. Taken together, these data confirm the hypothesis that cisplatin resistance increases proangiogenic capacity of A549 cells.

Angiogenesis is a cardinal feature of aggressive lung adenocarcinoma [63]. In addition to the canonical proangiogenic effects, increased VEGF expression has been correlated with cisplatin resistance in multiple carcinoma types, including lung adenocarcinoma [64, 65]. For example, cisplatin induction of VEGF upregulates microRNAs that promote autophagic cell survival in prostate cancer cells [66]. Increased proangiogenic capacity of Cis-R A549 spheroids in the current study (**Figure 5**) corresponded to an upregulation of VEGF-A mRNA (**Figure 4B**, P = 0.148). Suppression of angiogenesis or vascular normalization are therapeutic goals in oncology, and recent studies have suggested that cisplatin efficacy may be in part driven by antiangiogenic effects that augment tumor cell killing [67, 68]. More broadly, targeted anti-angiogenics are a common component of therapies for LC and other cancers [69]. While compounds such as bevacizumab (anti-VEGF-A monoclonal) for LC have shown statistical benefits in clinical trials, the significance of the effect on overall survival remains ambiguous [70, 71]. The impacts of tumor vascularization and anti-angiogenic therapy on the acquisition of drug resistance have been described [72–74], but it is unclear if the surviving drug-resistant populations of carcinoma cells are more proangiogenic. If so, the timing of anti-angiogenic therapy in the overall sequence of treatments in patients with recurrent disease could impact outcomes. Our results suggest that drug-resistant recurrent carcinomas could be more potently proangiogenic (**Figure 5E-J**). Therefore, the idea of reserving anti-angiogenics for second line combination therapies in the recurrence phase as a strategy to slow overall disease progression warrants further preclinical investigation.

### Conclusion

This proof-of-concept study demonstrated the ability to engineer features of increased aggressivity for MPS models of advanced cancer via selection for drug resistance in 2D culture (**Figure 1**). These features include: i) induction of pseudo-dormancy in persister cells defined by a markedly lower frequency of cell cycle participation (**Figure 2 and 3**), ii) mesenchymal-like transition (**Figure 2**), iii) dysmorphic architecture in 3D culture (**Figure 3**), iv) upregulation of progression-associated genes (**Figure 4**), and v) increased proangiogenic capacity (**Figure 5**). This study demonstrates the utility of using human cell-based MPS models to screen antiangiogenics in accordance with the increased NIH emphasis on new approach methodologies (NAMs) for preclinical research. Upon further validation, the observed association between a drug-resistant phenotype and increased proangiogenic capacity could suggest the relative importance of treatments targeting angiogenesis in the recurrence phase following first-line platinum-based therapies. Future work will focus on establishing cisplatin resistant derivatives of other carcinoma cell lines to investigate cell line-specific effects, derivatives of the same lines selected for resistance to other compounds to investigate compound-specific effects, and resistant derivatives of lung cancer patient-derived organoids to investigate the effects of individual variability on cisplatin resistance and the resultant phenotypes.

## Methods

### Cell Culture

Lung adenocarcinoma A549 cells [American Type Culture Collection (ATCC)] were cultured in Kaign’s F-12K medium (ATCC) supplemented with 10% fetal bovine serum (FBS, Corning). Normal human lung fibroblasts (HLFs, ATCC) were cultured in fibroblast basal medium with low serum kit (ATCC) and 2% fetal bovine serum (FBS). Human umbilical vein endothelial cells (HUVECs, ATCC) were cultured in vascular cell basal medium (ATCC) with endothelial cell growth kit-VEGF (ATCC). Early passage numbers (P2-P4) routinely tested for mycoplasma contamination were used for all cell types. A549 spheroids were cultured in 96 well round bottom ultra-low attachment plates (ThermoFisher 262162). Spheroids that were immunostained were seeded at 1000 cells per well and spheroids used in RT-qPCR experiments were seeded at 5000 cells per well. For RT-qPCR, 48 spheroids were collected per group for RNA isolation. Spheroids were cultured for 4 days.

Spheroids used in angiogenesis assays were seeded at 3,000 cells per well in elplasia 96 well microcavity plates (Corning 4442) and cultured for 3 days. One well was used per angiogenesis device. All culture media contained 1% antibiotic-antimycotic (Corning) and cultured in a humidified tissue culture incubator at 37°C and 5% CO_2_.

### Cisplatin Drug Response

Cisplatin (Selleckchem S1166) was dissolved in sterile de-ionized water according to manufacturer’s instructions and added to culture medium at 25μM. Once A549 cells reached confluency, cisplatin was administered twice at 48-hour intervals over a total period of 96 hours. The drug media was prepared fresh at every drug dose. After the 96-hour cisplatin exposure, media was changed to drug-free culture medium (F-12K with 10% FBS and 1% ABAM). Cells were passaged once, prior to being used in experiments.

### Crystal Violet

Cells were seeded at 1000 cells per well in a 96 well plate and cultured to confluency for 4 days. The media was replenished every 48 hours. The cells were treated with cisplatin at 0, 5μM, 15μM, 25μM, 35μM, 45μM, and 55μM for 96 hours. Following the 96-hour treatment regimen, cells were fixed in 4% paraformaldehyde for 15 minutes and washed with Dulbecco’s phosphate buffer solution (DPBS, Corning) prior to staining. Crystal violet (CV) (Sigma Aldrich V5265) solution was made at 1:10 in 10% methanol. The CV solution was added to the plate and incubated for 30 minutes at room temperature. Cells were washed with DPBS until the solution ran clear and set to dry overnight. Crystal violet was eluted with 33% acetic acid solution, and the absorbance was read on a SpectraMax i3x plate reader at 570nm.

### Flow Cytometry

Media was collected in the middle of cisplatin treatment to collect floating dead cells in suspension, as well as at end point. At end point, cells were washed with 1mL sterile PBS and then trypsinized with 0.05% trypsin-EDTA and incubated at 37°C for 6 minutes. Then trypsinization was blocked with 2mL media and then contents were collected for centrifugation at 1000 rotations per minute (rpm) for 5 minutes. Cells were resuspended in 2mL non-sterile PBS and then split into 2 flow cytometry tubes with 1mL each. One tube was used as an unstained control and another tube was stained with DAPI at a 1:1000 dilution. Samples were centrifuged at 1000 rpm for 5 minutes and then resuspended in 500uL non-sterile PBS. Flow cytometry was carried out using FACS Melody. Data was analyzed using FlowJo software. First forward scatter plot and side scatter plots were gated to exclude cell debris and doublets.

### MFOC Device Fabrication and Organ chip Seeding

A double membrane-free organ chip (MFOC) Polydimethylsiloxane (PDMS) device was fabricated using methods as described before [61, 62]. The double MFOC is composed of two rectangular central channels for gel loading, flanked by two media side channels. The device is assembled using 3 layers formed by PDMS soft lithography using 3D Stereolithography (SLA) printed molds. The bottom and middle layer combine to form the gel and media compartments. The top layer is a reservoir for media. Before gel injection, the organ chips were sterilized using Ultraviolet light in a biosafety cabinet for 20 min. The device is functionalized for ECM hydrogel anchorage using a 5 mg mL^−1^ polydopamine solution (PDA). Dopamine solution was prepared by mixing dopamine hydrochloride (Sigma-Aldrich) with 10 mm Tris-hydrochloride (Sigma–Aldrich) dissolved in ultra-pure water. Dopamine solution was sterile filtered using a 0.2 µM filter and injected central channels of double MFOC. Devices were incubated for 2 h at room temperature under ultraviolet irradiation, aspirated, and washed with ultra-pure water. PDA-treated devices were used within 1 week of coating.

The organ-chips were loaded using a protocol reported previously [62]. Briefly, HLF and HUVEC (2 × 106 cells mL−1 each) were admixed in 2.2 mg mL−1 collagen I (Corning), 5 mg mL−1 fibrinogen (Sigma-Aldrich), and 1 U mL−1 thrombin (Sigma–Aldrich). The cell-inoculated hydrogel precursor was injected into one of the central tissue lanes of the double MFOC devices and incubated for 15 min at 37 °C. The adjacent central tissue lane was seeded with one well of Cis-R or parent spheroids admixed in 50µL of 2.2 mg mL−1 collagen I (Corning), 5 mg mL−1 fibrinogen (Sigma–Aldrich), and 1 U mL−1 thrombin (Sigma–Aldrich) and incubated for 15 min at 37 °C. Control groups were loaded with the same hydrogel mixture without the spheroids. The outer side channels and media reservoirs with VEGF-free endothelial cell growth medium supplemented with 25 µg mL−1 aprotinin (EMD Millipore, 616370-20MG). The media was changed every 2 days. The devices were cultured for 10 days before fixation.

### Immunofluorescence Staining

**Cis-R cell monolayer staining:** Cell monolayers were seeded at 30,000 cells per well in 24 well plates and cultured for 4 days. Monolayers were fixed in 4% PFA for 15 minutes and washed with DPBS prior to immunostaining. Samples were blocked and permeabilized for 20 minutes in 1% bovine serum albumin (BSA) and 0.1% Triton-X, prepared in DPBS. Primary Ki-67 antibody (Rabbit polyclonal to Ki67 [ab15580], abcam), at 1:100 in 0.1% BSA prepared in DPBS and incubated at room temperature for 2 hours with gentle rocking. Secondary antibody (Alexafluor 594 goat anti rabbit IgG [150080], abcam), Hoescht 33342, and phalloidin was added at 1:500 in 0.1% BSA and incubated in the dark for 40 minutes with gentle rocking. Samples were washed with DPBS before confocal imaging.

**Cis-R Spheroid staining:** Spheroids were seeded at 1,000 cells/well and cultured for 4 days in 100uL F12k media supplemented with 10% FBS. Spheroids were fixed in 4% PFA for 45 minutes and washed with DPBS prior to immunostaining. Spheroids were blocked and permeabilized for 30 minutes in 1% BSA and 0.1% Triton-X, prepared in DPBS with gentle rocking. Primary Ki-67 antibody (Rabbit polyclonal to Ki67 [ab15580], abcam) at 1:100 in 0.1% BSA prepared in DPBS and incubated at room temperature for 2 hours with gentle rocking. Spheroids were washed with DPBS and gently rocked. Secondary antibody (Alexafluor 594 goat anti rabbit IgG [150080], abcam), Hoescht 33342, and phalloidin was added at 1:500 in 0.1% BSA prepared in DPBS and incubated in the dark for 1 hour with gentle rocking. Samples were washed with DPBS before confocal imaging.

**Vascular Staining in Devices:** Devices were fixed with 4% PFA for 1 hour at room temperature then refrigerated overnight. The following day devices were washed with DPBS three times for 15 minutes on the rocker prior to immunostaining. Devices were blocked and permeabilized in 1% BSA+ 0.2% Triton-X prepared in DPBS with Hoescht 33342 and phalloidin added at 1:250, and Ulex Europeas agglutinin I (UEA-1, Vectorlabs) added at 1:50. The side channels of the devices were loaded with 50uL and incubated at room temperature on the rocker for 1 hour and overnight in the fridge. The following day, devices were rocked at room temperature for one hour prior to being washed 5 times with DPBS for 3 hours.

### Confocal Imaging and Image Analysis

Samples were imaged in their respective plates using a Nikon Ti-2 Confocal Microscope at 10x objective on DAPI, FITC, and TRITC channels. Spheroids were imaged by creating a z-stack using a 10 µm step size for 20 steps. The FITC channel was used as a guide to determine the top and bottom of the spheroid. Stained angiogenesis tissues were fixed in position on a glass slide and imaged on an inverted Nikon C2 laser scanning confocal microscope (LSCM) equipped with a Nikon DS-FI3 camera. A large composite 6×1 stitched image was acquired from each sample of sprouting angiogenesis assay to ensure convergence of morphological variables. Max intensity projection of Z-stacks of the samples were exported as TIFFs for morphometric analysis. All images were denoised prior to image analysis. **2D cell morphology:** Morphology of 2D images was conducted using an open-source CellProfiler project pipeline (Broad Institute, Cambridge, MA, USA). Nuclei area, nuclei diameter, cell area, and cell diameter were assessed. Ki67 analysis was assessed by counting cell nuclei and Ki67 positive overlaps. The final Ki67 index was given as a ratio of the total Ki67 count divided by total nuclei count. **Spheroids Morphology:** Image analysis was conducted using ImageJ software. Channels were separated and max z-projections were created. The DAPI and FITC channels were then thresholded and made binary. Area, circularity, roundness, and solidity was computed using the measure particle plugin.

**Sprout Angiogenesis Morphometry:** All Image analysis was completed in MATLAB (R2021b). Vascular network images were smoothed using an edge-preserving filter with a Gaussian kernel, and a threshold was applied to remove remaining low-intensity noise. We used a pretrained deep neural network to denoise each image and adaptive histogram equalization was used to standardize contrast across the image set. We then segmented pre-processed images and quantified morphometric parameters using an open-source automated segmentation tool (REAVER). For quantification of angiogenic sprout growth, differential interference contrast (DIC) images of the central channels of each chip were used to identify the boundary between the TME compartment and vascular margin, and morphometric quantification was performed on the regions bound by the central boundary and the ends of the TME compartment.

### RT-qPCR

Cell pellets, spheroids, and tissues were stored at 4°C in RNALater solution (Invitrogen AM7020). RNA extraction was performed using the Qiagen’s RNeasy kit (74104) according to the manufacturer’s protocol. RNA purity and concentration was measured using a nanodrop spectrophotometer. Samples with 260/280 and 260/230 outside 2.0 ± 0.3 were discarded. RNA samples were stored in −80°C. For cDNA synthesis, 1 μg of RNA was used with qScript cDNA supermix (Quantabio, 95048-100) according to manufacturer’s instructions. Reaction was carried out in a thermocycler using the following conditions: 25°C for 5 min; 42°C for 30 min; 85°C for 5 min; 4°C hold. Following cDNA synthesis reaction, samples were diluted 1:10 with nuclease-free water and stored at −20°C. Working primers were prepared at 1:10 by adding 10μL of both the forward and reverse DNA Oligo into 180μL nuclease-free water and stored at 4°C. RT-qPCR was completed on the Applied Biosystems Step One Plus instrument and was quantified using Power SYBR green master mix (Applied Biosystems, 4368577). The following recipe was used for each reaction with a total volume of 20μL: 10μL Power SYBR green master mix, 1 μL working primer, 6 μL nuclease-free water, and 3μL cDNA. Gene expression levels were normalized to housekeeping control gene GAPDH and assessed using the 2^−ΔΔCt^ method.

The following primer sequences were used:

GAPDH Forward (TTAAAAGCAGCCCTGGTGAC) Reverse (TCTGCTCCTCCTGTTCGAC)

VEGFA Forward (CGAGGGCCTGGAGTGTGT) Reverse (CCGCATAATCTGCATGGTGAT)

IL8 Forward (GGTGCAGTTTGCCAAGGAG) Reverse (TTCCTTGGGGTCCAGACAGA)

CDH1 Forward (AGGTGACAGAGCCTCTGGATAGA) Reverse (TGGATGACACAGCGTGAGAGA)

αSMA Forward (CCGACCGAATGCAGAAGGA) Reverse (ACAGAGTATTTGCGCTCCGAA)

TGFβ1 Forward (TACCTGAACCCGTGTTGCTCTC) Reverse (GTTGCTGAGGTATCGCCAGGAA)

EGFR1 Forward (AGGCACGAGTAACAAGCTCAC) Reverse (ATGAGGACATAACCAGCCACC)

PDL1 Forward (CTGGCATTTGCTGAACGCAT) Reverse (AGGTCTTCCTCTCCATGCAC)

MMP2 Forward (TTTGAGTCCGGTGGACGATG) Reverse (GCTCCTCAAAGACCGAGTCC)

### Statistical Analysis

All experiments if not stated were repeated n = 3 times. Images were preprocessed and segmented per the above image analysis methodology. All statistical analyses were completed in GraphPad Prism V 10.3.1 (GraphPad Software Inc.). Comparison between two experimental groups were analyzed using an unpaired t-test. Welch’s correction was used if the standard deviation between the two groups was significantly different (p<0.05). Comparison between more than two experimental groups were analyzed using a one-way ANOVA Dunnett’s multiple comparison if not stated. The data are presented as mean ± SEM. Sample sizes and the number of independent experiments performed are indicated in the respective figure captions. Statistical significance was assigned as not significant (ns) when p > 0.05; or significant with *p ≤ 0.05; **p ≤ 0.01, ***p ≤ 0.001, ****p ≤ 0.0001.

## Disclosures

**Clinical trial registration:** N/A.

**Ethical approval:** N/A.

**Informed consent:** N/A.

## Author contributions

E.A.O., G.W.K., and O.M.K.A. contributed to experimental design, established methods, and conducted the experiments. M.J.M. conceived the study, directed the project, and wrote the paper. G.W.K. helped prepare and edit the paper.

## Data Availability

The data that support the findings of this study are available from the corresponding author upon reasonable request.

## Supplemental Figures

**Supplemental Figure 1:**
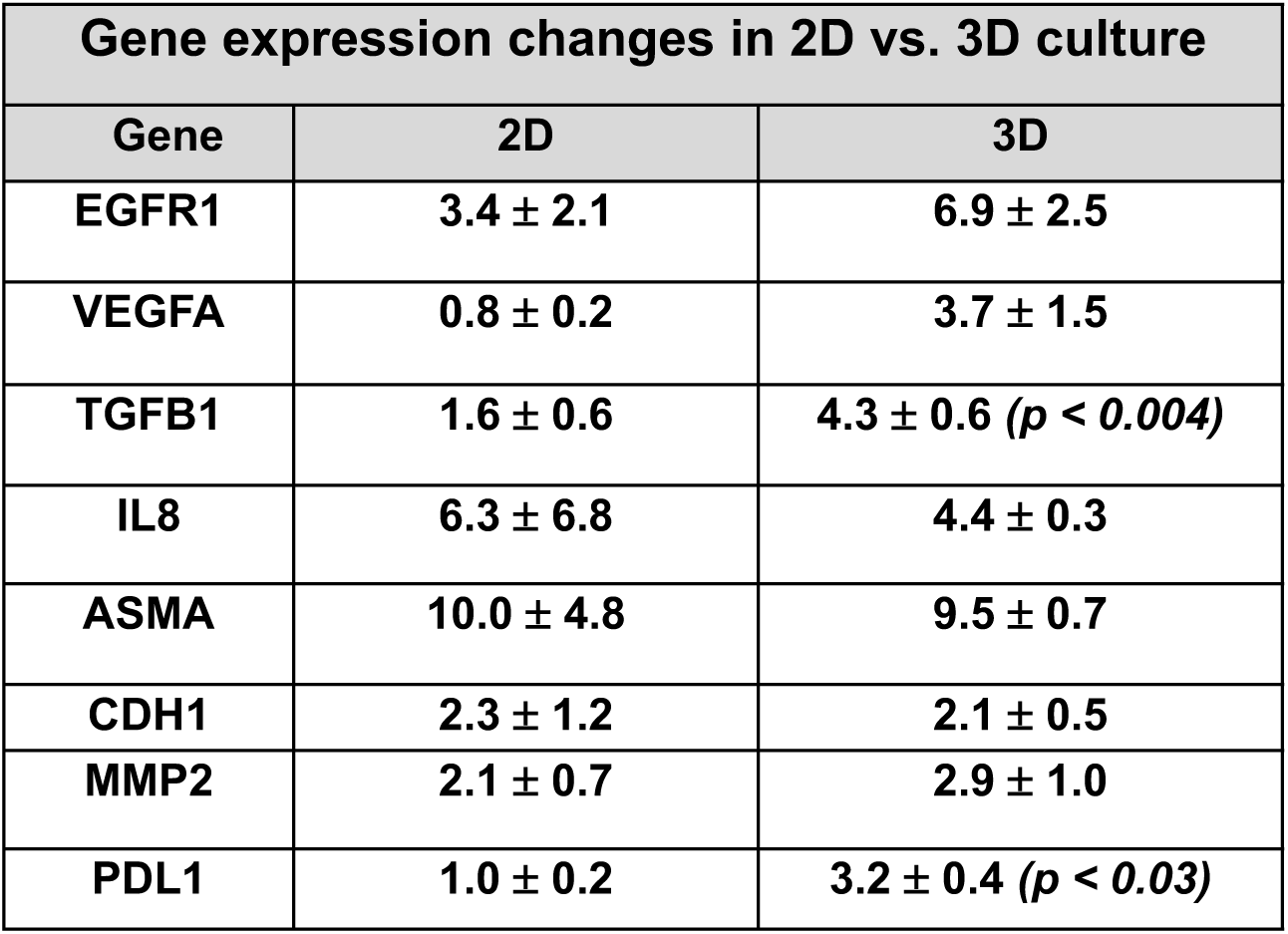
Table showing the relative mRNA fold changes between 2D and 3D cisplatin-resistant A549 lung adenocarcinoma tissues. Means ± standard deviation denotes the mRNA expressions relative to their parent A549 in 2D and 3D control groups. N= 3 independent experiments. Statistical analysis between 2D and 3D groups was performed using an unpaired t-test with Welch’s correction in GraphPad Prism v10.3.1. Statistical significance is indicated by * = p < 0.05, ** = p < 0.01, *** = p < 0.001. Only significant differences were denoted.

**Supplemental Figure 2:**
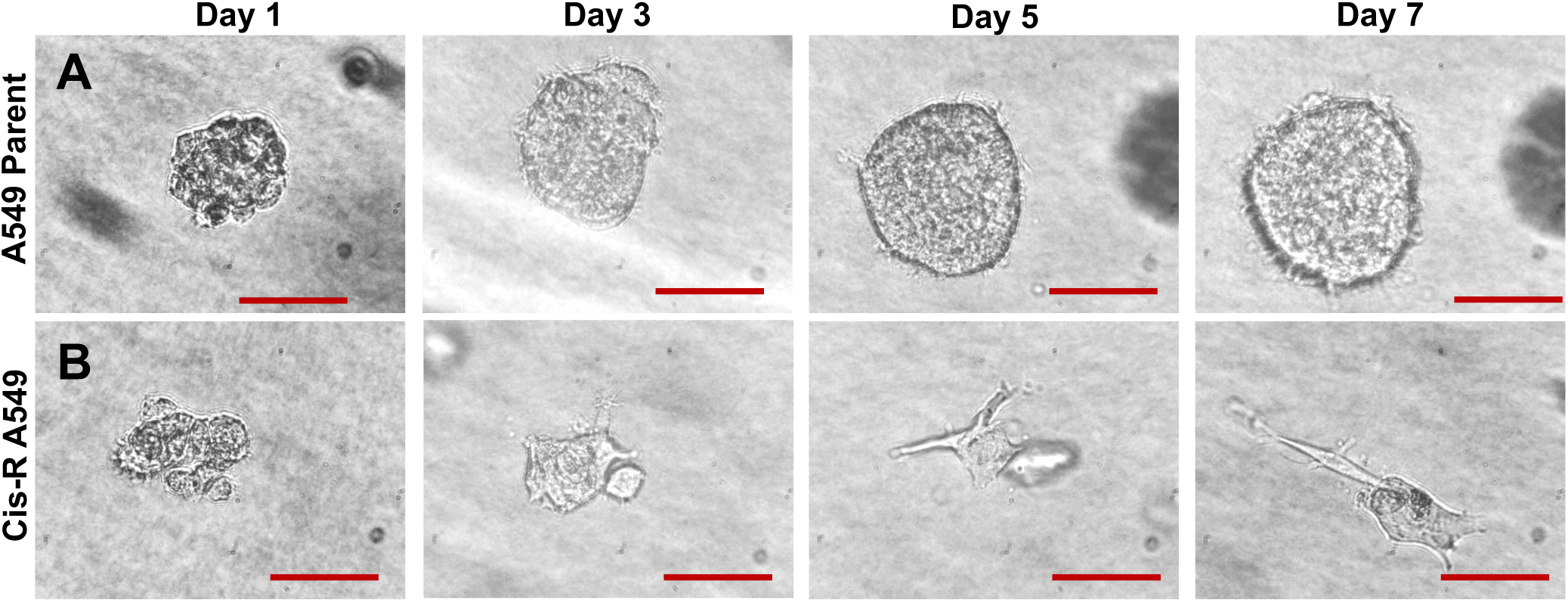
Time course images of parent A549 and cisplatin-resistant spheroids cultured in 2.5 mgmL^−1^collagen hydrogels for 7 days. Scale bar = 200um. A: Representative brightfield images of parent A549 spheroids taken on day 1, day 3, day 5, and day 7. B: Representative brightfield images of CisR A549 spheroids taken on day 1, day 3, day 5, and day 7.

